# CTG clade-specific proteins of the RSC chromatin remodeling complex regulate cell cycle progression of a critical priority fungal pathogen, *Candida albicans*

**DOI:** 10.1101/2025.10.06.680682

**Authors:** Ankita Joshi, Gayatri Brahmandam, Harini Kannan, Shilajit Roy, Sandhya Subramanian, Amartya Sanyal, Santanu Kumar Ghosh

**Affiliations:** Department of Biosciences and Bioengineering, Indian Institute of Technology Bombay, Powai, Mumbai, India; Department of Biological Sciences, Birla Institute of Technology and Science, Pilani, Hyderabad Campus, Jawahar Nagar, Kapra Mandal, Medchal District, Telangana, India

**Keywords:** RSC complex, chromatin remodeling, fungal pathogen, *Candida albicans*, cell cycle, virulence

## Abstract

The RSC and the homologous chromatin remodeling complexes are known to regulate cell cycle progression in various organisms, including *Saccharomyces cerevisiae*, *Drosophila*, and *Homo sapiens*. In this work, we characterized the role of two novel CTG clade-specific proteins (Nri1 and Nri2) of the RSC complex in the regulation of cell cycle progression in a critical priority fungal pathogen, *Candida albicans*. We observed that Nri1, alone or along with Nri2, regulates cell cycle progression at multiple stages. The *nri1Δ/Δ* and *nri1Δ/Δ nri2Δ/Δ* mutants exhibited transient cell cycle arrest, defective spindle morphology, and cytokinesis. Transcriptomic analysis supported these mutant phenotypes and indicated a broad role of Nri proteins in the cell cycle. From our results, we conclude that Nri proteins are crucial for *C. albicans* proliferation and fitness.

**Importance:** The composition of the essential RSC chromatin remodeling complex exhibits species-specific divergence, harboring unique subunits with distinct functions. In this study, we report that two fungal CTG clade-specific proteins of the *C. albicans* RSC complex, namely Nri1 and Nri2 can promote *C. albicans* fitness through regulating its cell cycle progression at multiple stages. Fitness defect along with stressor sensitivity and differential expression of the genes regulating pathogenesis in the *nri* mutants indicate potentiality of the Nri proteins as anti-*Candida* drug targets.

## Introduction

*Candida albicans*, categorized as a critical priority fungal pathogen by World Health Organization, inhabits various niches in human host as a harmless commensal (Brown et al. 2012). It is present in the oral cavity, gastrointestinal and genito-urinal tract, and on the skin of majority of the population (Ghannoum et al. 2010; Drell et al. 2013; Ng et al. 2015; Silva Dantas et al. 2016; Nash et al. 2017). However, in case of compromised host immunity and disbalanced microbiota due to various reasons, it can cause mucosal infections or life threatening systemic infection (Gudlaugsson et al. 2003; Kim and Sudbery 2011; Poulain 2015; Pappas et al. 2018). *C. albicans* uses a diverse range of strategies to improve its survival in the host niches, ultimately increasing virulence. It is a polymorphic fungus, can adhere and invade host mucosa, form biofilms, generate ‘beneficial aneuploidy’ and has the potential to develop resistance against antifungal drugs (Mayer et al. 2013; Alves et al. 2020; d’Enfert et al. 2021). Most of these pathogenic attributes fundamentally rely on transcriptional regulation, where modulation of chromatin on demand plays a crucial role. Consequently, the chromatin factors such as histone-modifying enzymes and ATP-dependent chromatin remodeling complexes have emerging roles in fungal pathogenesis (Rosa and Kaufman 2012; Balachandra and Ghosh 2022). Histone modifications impart biological functions through *cis*-effect on the chromatin itself and through *trans*-effect, they recruit various ‘reader proteins’ including chromatin-remodeling complexes to alter the chromatin accessibility by repositioning, exchanging, or removing the nucleosomes (Clapier and Cairns 2009; Längst and Manelyte 2015).

Various reports highlight the importance of chromatin factors in *C. albicans* biology. Histone acetylase Rtt109 is crucial for nucleosome assembly in the S phase, and *C. albicans* lacking *RTT109* exhibits hypersensitivity to DNA-damaging agent hydroxyurea with activation of DNA damage response (Lopes et al. 2010). *C. albicans* exhibits defective spindle morphology and anaphase progression when histone deacetylases Hst3 activity is inhibited (Conte et al. 2022). In this organism, Sir2 histone deacetylase is also reported to regulate rDNA stability and mitotic exit (Price et al. 2019). Besides regulating cell cycle progression, the histone modifiers are also known to influence cell wall integrity, morphological transitions, genome stability, stress response, and virulence in *C. albicans* (Raman et al. 2006; Lopes et al. 2010; Tscherner et al. 2015; Shivarathri et al. 2019; Conte et al. 2022; Yang et al. 2024). Exchange of canonical histones with timely deposition of histone variants H3V^CTG^ and H2A.Z was also shown to contribute to morphological transitions and biofilm formation in *C. albicans* (Rai et al. 2019; Qasim et al. 2021; Brahma et al. 2023). Similarly, the chromatin remodeling complexes also regulate several crucial cellular functions in various organisms. SWI/SNF superfamily of remodelers, predominantly the RSC chromatin remodelling complex, controls transcriptional activation and repression (Laurent et al. 1993; Soutourina et al. 2006; Balachandra et al. 2020), mitotic progression (Cao et al. 1997; Prasad et al. 2019), DNA damage repair (Chai et al. 2005; Shim et al. 2007; Czaja et al. 2014), cytoskeletal organization (Chai et al. 2002), kinetochore clustering (Prasad et al. 2019), cohesion (Huang et al. 2004; Muñoz et al. 2019; Prasad et al. 2019; Muñoz et al. 2022), and chromosome segregation (Hsu et al. 2003) in fungal organisms.

Earlier, through mass spectrometry-based identification of the *C. albicans* RSC chromatin remodeling complex, we discovered the presence of two novel CTG-clade specific subunits, namely Nri1 and Nri2 (Balachandra et al. 2020). Given the key roles of several core subunits of the RSC complex in *C. albicans* proliferation and pathogenicity (Prasad et al. 2019; Balachandra et al. 2020; Prasad 2022; Prasad et al. 2022; Prasad 2024), we hypothesized that CTG-clade specific Nri proteins would also significantly influence *C. albicans* biology and have the potential to be ideal drug targets. In this work, we demonstrate that the *nri* mutants are defective in cell cycle progression. Moreover, they show a kinetochore organization defect, impaired spindle morphology, and altered chromatin compaction. RNA sequencing (RNA-seq) analysis of *nri* mutants revealed differential expression of several genes involved in those processes, supporting the observed phenotypes. Altogether, we conclude that the Nri proteins indeed can regulate a broad range of cellular processes. Rapid adaptation through proficient stress response, increased drug resistance cases, and limited antifungal treatment options due to associated host toxicity (Pristov and Ghannoum 2019) advocate the dire need for developing novel therapeutic targets. Chromatin-remodeling proteins, being regulator of fungal fitness by means of diverse pathways, can in fact limit both fungal growth and adaptation, and hence, the virulence. In this context, our work postulates the role of fungal CTG clade-specific RSC complex proteins in regulating *C. albicans* fitness, highlighting their importance as therapeutic targets.

## Results

### *nri1Δ/Δ nri2Δ/Δ* double mutant is synthetically sick indicating genetic interaction between the *NRI* genes

We reported earlier that the *nri1Δ/Δ* mutant had growth defect under standard growth conditions, and it was hypersensitive to various stress-inducing agents mimicking physiologically relevant hostile conditions; on the other hand, *nri2Δ/Δ* mutant only displayed thermosensitivity (Balachandra et al. 2020). To determine the impact of combined loss of *NRI1* and *NRI2* genes on *C. albicans*, we constructed a double mutant of these genes. Similar results were obtained for *nri1Δ/Δ* and *nri2Δ/Δ* single mutants as reported in Balachandra et al., 2020 (Figure 1A, Figure S1A), whereas the *nri1Δ/Δ nri2Δ/Δ* double mutant exhibited a severe growth defect under standard growth conditions (Figure 1A, B). The doubling time for *nri1Δ/Δ* and *nri1Δ/Δ nri2Δ/Δ*, calculated based on the growth curve, were 104.6 min and 121.6 min, respectively, which were significantly higher than that of the wild-type (WT) strain that took 81.84 min to double (Figure 1C). The observed synthetic sick phenotype of the *nri1Δ/Δ nri2Δ/Δ* double mutant confirmed a genetic interaction between the *NRI* genes. As expected, the *nri1Δ/Δ nri2Δ/Δ* double mutant also displayed hypersensitivity to various physiologically relevant stress conditions, similar to the *nri1Δ/Δ* mutant (Figure S1B). As *nri2Δ/Δ* single mutant displayed no major phenotypes (except thermosensitivity as reported earlier), we then focused on *nri1Δ/Δ* single mutant and *nri1Δ/Δ nri2Δ/Δ* double mutant for further characterization.

**Figure 1:**
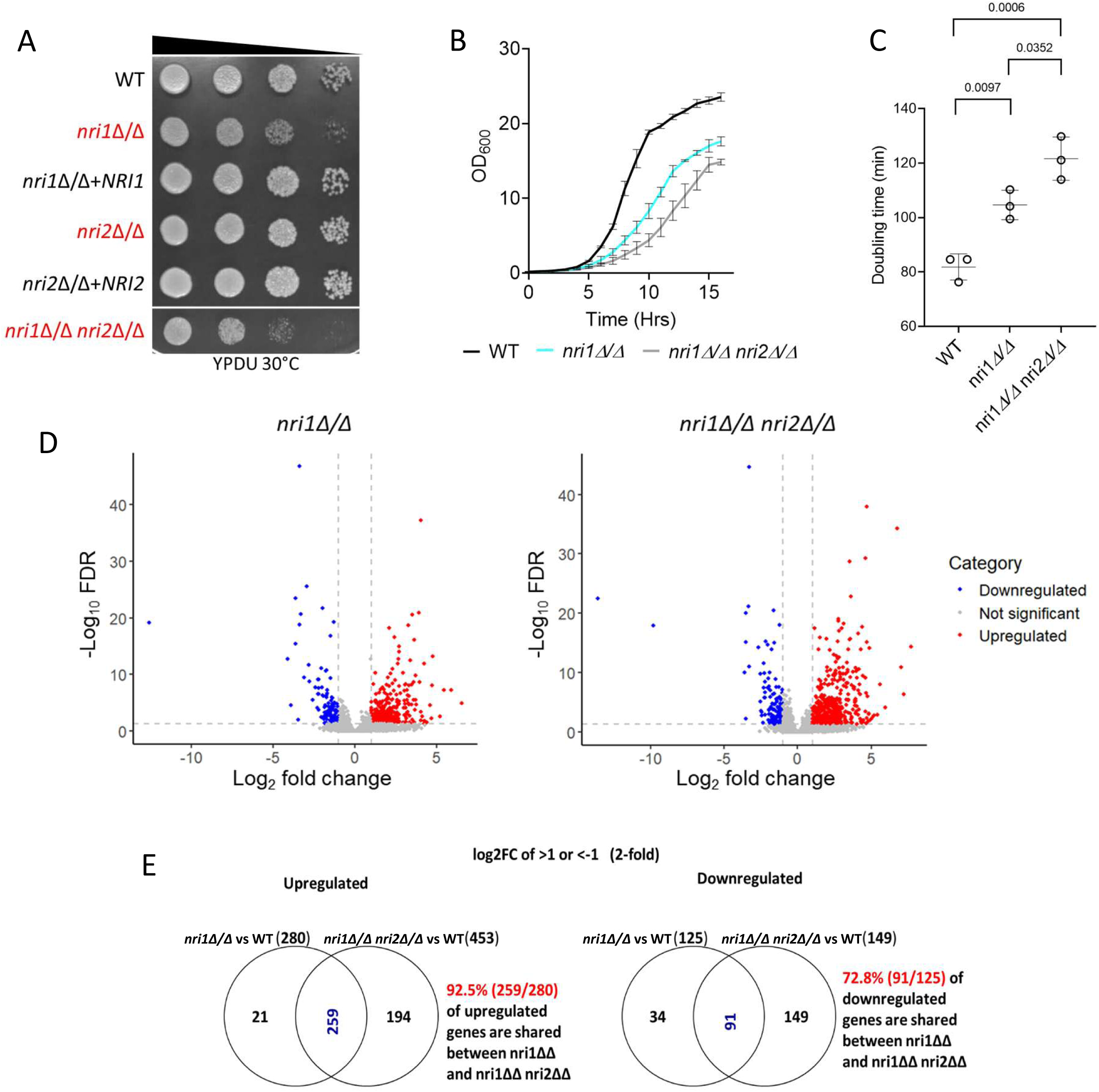
*nri1*Δ/Δ *nri2*Δ/Δ double mutant shows synthetic sick phenotype. **A.** 10-fold serially diluted cells of the indicated strains were spotted on the YPDU plate and incubated at 30 °C for 48 hrs and imaged. **B.** The growth curve was measured for WT, *nri1*Δ/Δ mutant, and *nri1*Δ/Δ *nri2*Δ/Δ double mutant strains at standard growth conditions. OD_600_ was recorded every 60 mins. Mean values with standard deviation from three independent experiments were plotted. **C.** Doubling time of the strains was calculated from the growth curve. Data from three biological replicates are plotted. Error bars indicate standard deviation. **D.** Volcano plot representations of the differentially expressed genes (DEGs) for *nri1Δ/Δ* w.r.t. WT (*nri1Δ/Δ*) and *nri1Δ/Δ nri2Δ/Δ* w.r.t. WT (*nri1Δ/Δ nri2Δ/Δ*) samples. Significantly altered genes with the threshold of - 1<log_2_(Fold Change)>1 and FDR (adjusted p-value) <0.05 are highlighted in blue and red colours, respectively. **E.** Venn diagram indicating overlap between the upregulated and downregulated genes in *nri1Δ/Δ* and *nri1Δ/Δ nri2Δ/Δ* mutants.

### Absence of Nri proteins causes global alterations in the transcriptome profile

As Nri proteins are a part of the RSC chromatin remodeling complex that has myriad functions (Prasad et al. 2019; Balachandra et al. 2020), high-throughput RNA-seq analysis was performed to understand the impact of the absence of *NRI1* and *NRI2* genes on the *C. albicans* transcriptome profile. With the absolute fold-change of 2 and FDR threshold of <0.05 (adjusted p-value), the absence of *NRI1* gene resulted in 280 up- and 125 down-regulated transcripts, accounting for 6.26% (405/6468) of the total transcripts in haplotype A of *C. albicans* strain SC5314 assembly 22. The combined loss of *NRI1* and *NRI2* genes had a relatively greater impact on transcriptomic changes with 453 up- and 149 down-regulated transcripts, comprising 9.3% (602/6468) of the total transcripts, corroborating genetic interaction between the two genes (Figure 1D, E). Gene ontology (GO) analysis was done using GO slim mapper tool of the Candida Genome Database (CGD, www.candidagenome.org), which indicated that the differentially expressed genes (DEGs) regulated various growth or fitness related processes such as “transport”, “translation”, “organelle organization”, “protein catabolic process”, “RNA metabolic process”, “cell cycle”, “cellular homeostasis”, “ribosome biogenesis”, and “cytoskeleton organization”. Apart from this, DEGs were also from the processes that influence virulence of the organism, such as “response to stress”, “filamentous growth”, “interspecies interaction”, “cell wall organization”, “biofilm formation”, and “cell adhesion” (Figure 2A). This analysis indicates that the Nri proteins regulate both growth and virulence attributes. To pinpoint the genes misregulated in the mutants, detailed analysis of the DEGs revealed a significant downregulation of *APC11* transcript, an ortholog of the anaphase-promoting complex component, in both *nri1Δ/Δ* and *nri1Δ/Δ nri2Δ/Δ* mutants compared to WT. DNA replication regulating genes *RNR3*, *CDC45*, and *ORC1*, transcription regulatory gene *KNS1*, kinetochore protein encoding genes *NSL1* and *DAD4*, and mediator complex subunit encoding gene *MED9* were also found to be differentially expressed in the *nri1Δ/Δ nri2Δ/Δ* mutant. Out of these DEGs, *CDC45* and *ORC1* also showed significant downregulation in *nri1Δ/Δ* single mutant. It is worth mentioning that downregulation of *CDC45*, *ORC1*, *NSL1*, and *MED9* was observed at fold change cutoff of 1.5-fold, a commonly used parameter in several studies (Linde et al. 2015; Balachandra et al. 2020; Kumwenda et al. 2022; Shivarathri et al. 2024). Several oxidative stress-related genes were differentially expressed in both mutants, supporting the observed H_2_O_2_ sensitivity (Figure S1B). Several transcription factors and epigenetic regulators, such as *RON1*, *ZCF25*, *ZCF26*, *HIR1*, etc. exhibited differential expression, suggesting the possible indirect regulatory roles of *Nri1* and *Nri2* proteins (Figure 2B).

**Figure 2:**
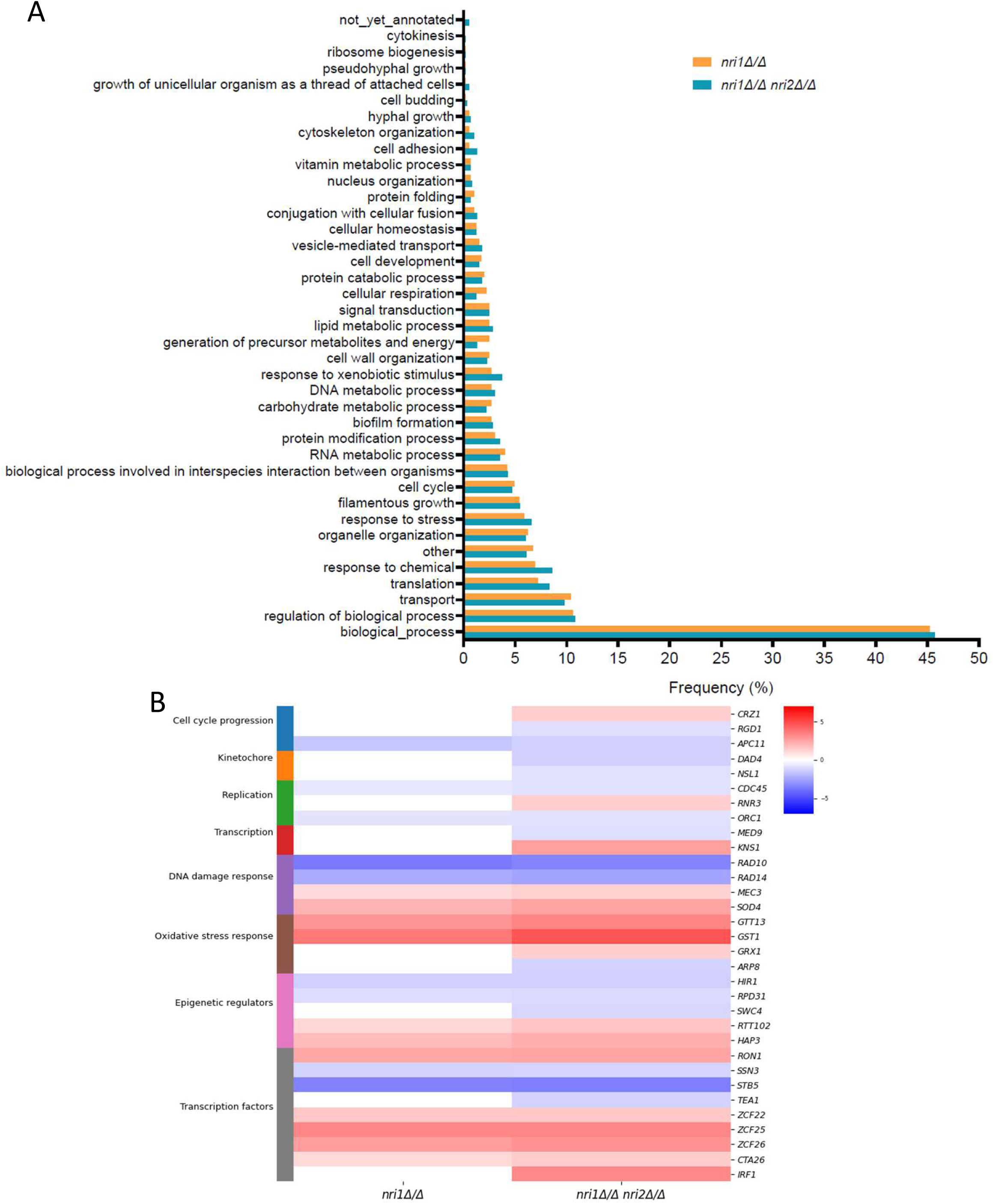
*nri* mutants exhibit global alterations in the transcriptome profile. **A.** Gene Ontology (GO) analysis of DEGs in *nri1Δ/Δ* w.r.t. WT (*nri1Δ/Δ*) and *nri1Δ/Δ nri2Δ/Δ* w.r.t. WT (*nri1Δ/Δ nri2Δ/Δ*) samples. Y-axis indicates various GO terms of the biological processes and X-axis indicated percentage of the DEGs encompassing that GO term. **B.** Heat map of DEGs in the indicated samples in various processes relevant to cell proliferation and virulence. Log_2_ fold change for the DEGs with FDR (adjusted p-

### *nri* mutants exhibit altered cell cycle progression

Due to the observed growth defect and DEGs under “cell cycle”, we first determined the cell death frequency in the mutants by staining the log-phase cells with propidium iodide, but observed no significant defects in the mutants (Figure S1C). We then analyzed the budding index in the asynchronously growing log-phase cells to understand the defects the mutants may harbor while progressing through the cell cycle. The behavior of the DAPI-stained nucleus with respect to the bud morphology was used for determining the cell cycle stages (Figure 3A). We did not observe any defects in the gross nuclear morphology and segregation in the mutants (Figure 3A, S1D). However, both the mutants showed a significant increase in the percentage of multi-budded cells, 14.1% in *nri1Δ/Δ* mutant and 25.5% in *nri1Δ/Δ nri2Δ/Δ* double mutant as compared to only 4.5% in WT strain (Figure 3A). Multi-budded cells might arise from defective cytokinesis/mitotic exit or cell separation defect. To address this, we treated the cells using zymolyase, which removes the cell wall and thus separates the cells having separation defects. However, the treatment could not change the percentage of multi-budded cells, indicating that the multi-budded cells exhibited cytokinesis defect rather than cell separation defect (Figure S2).

**Figure 3:**
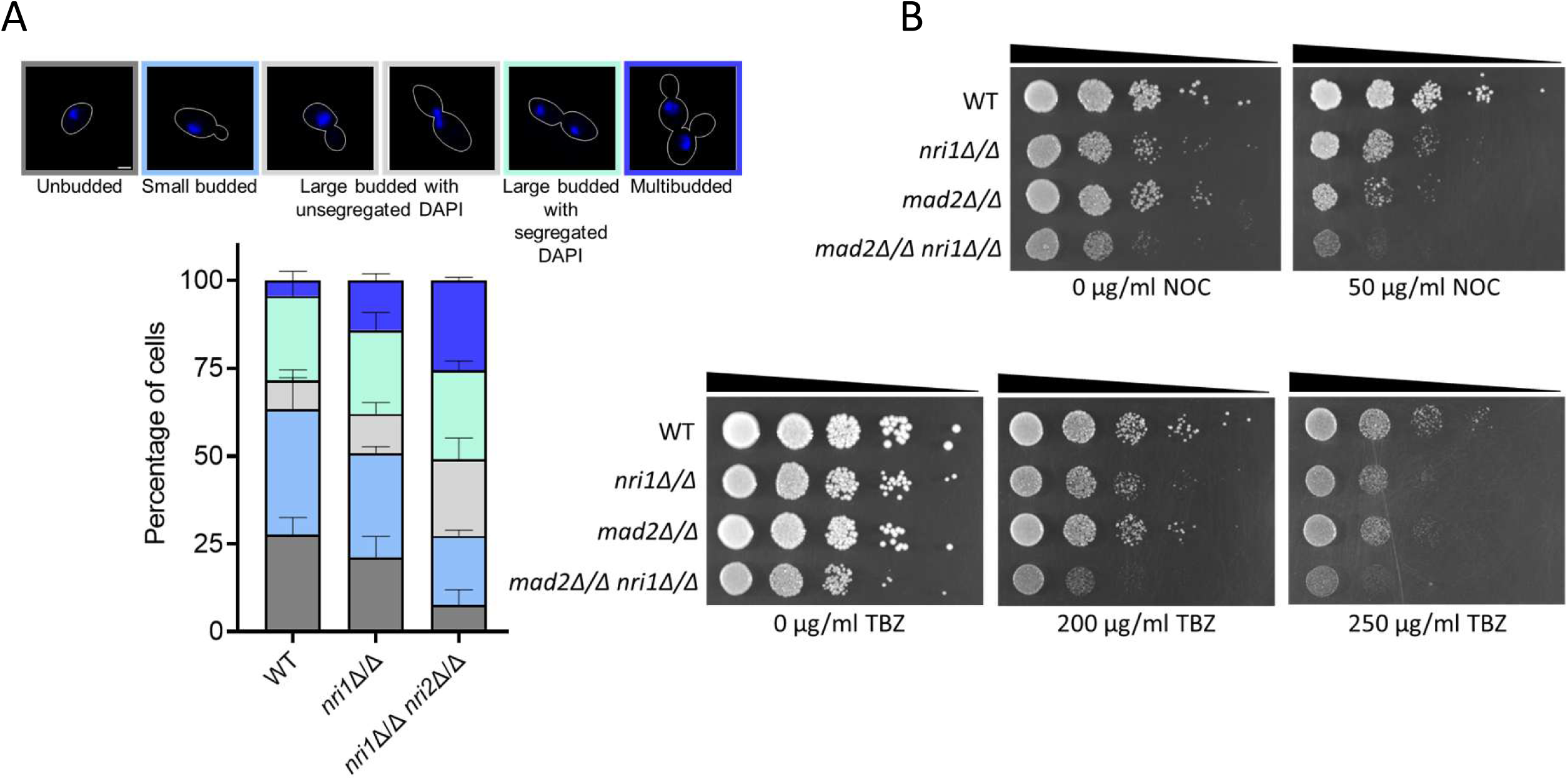
*nri* mutants display defective cell cycle progression. **A.** Above, representative images of different cell morphologies across the cell cycle stages based on bud size and nuclear localization. Scale bar 2 µm. Below, stacked bar graph showing the quantitation of the cell morphologies. Error bars indicates standard deviation. Data obtained from three biological replicates. N = 300. Statistical analysis performed by one-way ANOVA. WT, wild type. **B.** NOC and TBZ sensitivity was analyzed by spotting assay. 10-fold serial dilutions of the cells of the indicated strains were spotted on YPDU plates containing indicated concentrations of NOC and TBZ. Plates were incubated at 30°C for 48 hrs and imaged.

Notably, a significantly higher percentage of *nri1Δ/Δ nri2Δ/Δ* cells also showed large buds with unsegregated DAPI (21.8%), a phenotype similar to RSC mutants (Hsu et al. 2003; Wang and Cheng 2012; Prasad et al. 2019), indicating possible arrest at G2/M stage (Figure 3A). To understand whether this arrest is mediated by the activation of the spindle assembly checkpoint (SAC), *MAD2*, a SAC gene (Li and Murray 1991), was deleted in *nri1Δ/Δ* mutant. We observed synthetic sick phenotype of the *mad2Δ/Δ nri1Δ/Δ* double mutant as compared to their individual single mutants in the presence of microtubule depolymerizing drugs, nocodazole and thiabendazole. The double mutant showed a slight reduction in the growth rate in the absence of the drugs as well, altogether indicating a genetic interaction between *MAD2* and *NRI1* genes (Figure 3B).

### *nri* mutants display defects in spindle morphology at the later stage of the cell cycle

The cell cycle progression defect and genetic interaction with the *mad2Δ/Δ* mutant indicates that microtubule or kinetochore related defects may exist in the *nri* mutants. Since abnormal spindle morphologies were observed in Sth1 depletion mutant in *C. albicans* (Prasad et al. 2019), we therefore tested the spindle morphology in the *nri* mutants by live cell imaging of the asynchronously growing log-phase cells harbouring Cse4-GFP and Tub1-mCherry. Cells were categorized in two groups based on the cell cycle stages, namely pre-anaphase and post-anaphase judged by the bud size and Cse4-GFP signal, respectively. For all the tested strains, we did not observe any significant defect in the spindle morphology in pre-anaphase cells having no, small or large bud (Figure 4A). The post-anaphase cell population was further categorized into early anaphase and late anaphase based on pole-to-pole length which was judged by Cse4-Cse4 distance (4-6 µm as early anaphase and > 6 µm as late anaphase, respectively). We observed a significant increase in the cells with abnormal spindles in both early and late anaphase in *nri1Δ/Δ nri2Δ/Δ* double mutant and only in late anaphase in *nri1Δ/Δ* mutant (Figure 4B). Despite presence of abnormal spindles, no significant difference in pole-to-pole length was observed between wild type and the mutants (Figure 4C).

**Figure 4:**
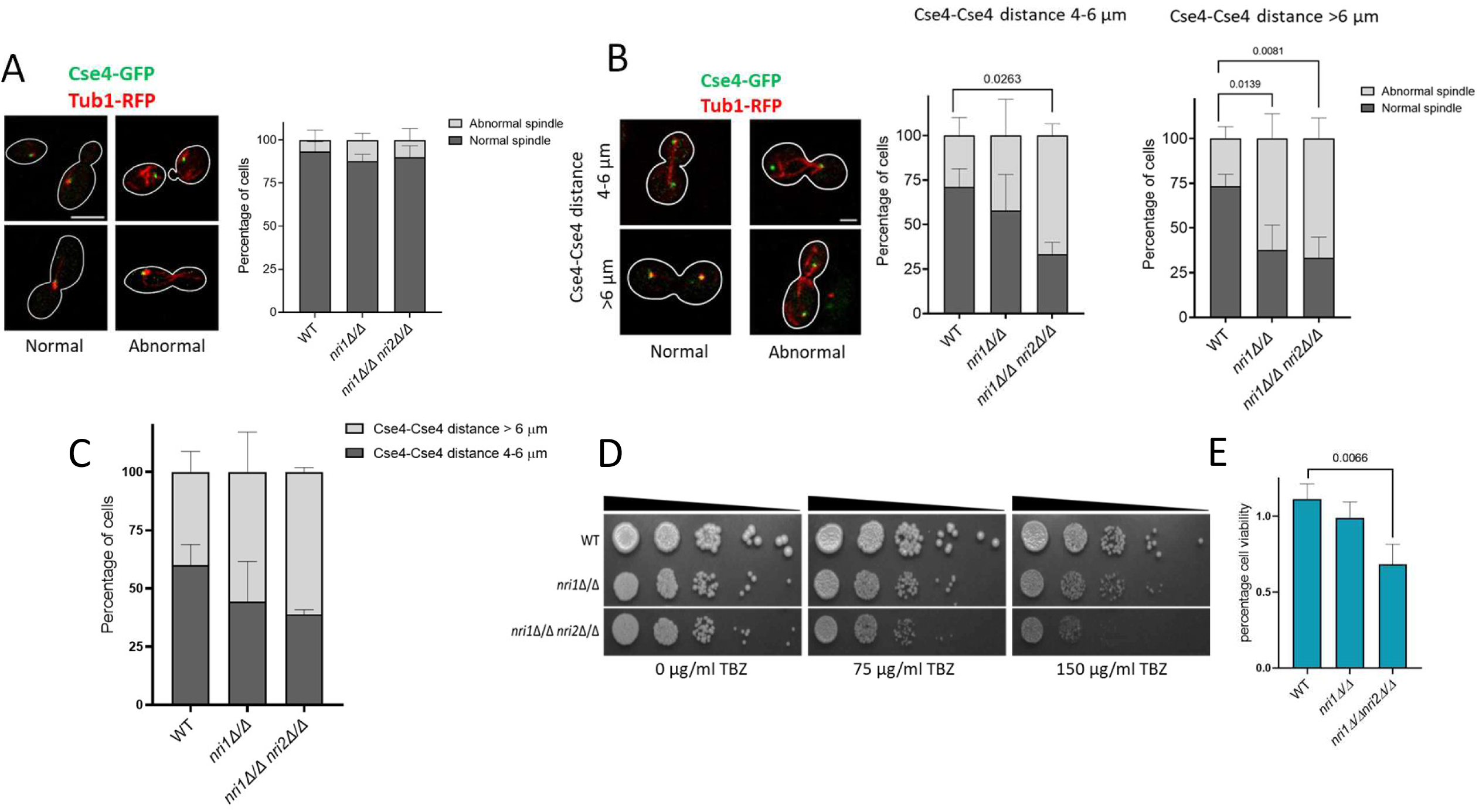
*nri* mutants exhibit abnormal spindle morphology at the later stage of the cell cycle. **A.** Left, representative images of normal and abnormal spindle morphologies in the pre-anaphase cells with no and small bud (top) and large bud (bottom). Scale bar 2 µm. Right, stacked bar graph of the percentage of pre-anaphase cells was plotted with the data from three independent experiments. Error bar indicates standard deviation. Statistical analysis was done by one-way ANOVA. N = 90. **B.** Left, representative images of normal and abnormal spindle morphologies in post-anaphase cells categorized into two groups based on the Cse4-Cse4 distance. Scale bar 2 µm. Middle and right, stacked bar graphs of the percentage of cells was plotted for both the groups with the data from three independent experiments. Error bar indicates standard deviation. Statistical analysis was done by one-way ANOVA. N = 45. **C.** Stacked bar graph of the percentage of post-anaphase cells harboring indicated Cse4-Cse4 distance was plotted with the data from three independent experiments. Error bar indicates standard deviation. Statistical analysis done by one-way ANOVA. N = 90. **D.** 10-fold serial dilutions of the cells of the indicated strains were spotted on YPDU plates containing indicated concentrations of TBZ. Plates were incubated at 30°C for 48 hrs and imaged. **E.** Bar graph showing the percentage of cell viability of the indicated strains in presence of 150 µg/ml TBZ from three biological replicates. Error bars indicate standard deviation. Statistical analysis was done by one-way ANOVA. Only significant p-value are mentioned in the graphs.

Cells with abnormal kinetochore or microtubule functions exhibit sensitivity to antimitotic drugs (Hyland et al. 1999; Ghosh et al. 2001; Anbalagan et al. 2024). As we observed spindle defects, and cell cycle arrest in the *nri* mutants, we tested the sensitivity of these mutants to thiabendazole (TBZ), an antimitotic microtubule-depolymerizing drug. Cells were spot inoculated on the plates containing 0, 75 or 150 µg/ml TBZ. *nri1Δ/Δ nri2Δ/Δ* double mutant showed hypersensitivity at both the concentrations (Figure 4D). CFU counting was done to quantify the viability of the mutants at 150 ug/ml TBZ concentration. We observed a significant drop in viability of *nri1Δ/Δ nri2Δ/Δ* double mutant (Figure 4E). However, *nri1Δ/Δ* single mutant only showed marginal sensitivity and viability loss at the tested concentrations (Figure 4D, E) which corroborates with a lesser spindle morphology defect in the single mutant compared to the double mutant.

Previous reports suggest a correlation between KT and spindle integrity (Hofmann et al. 1998; Jones et al. 2001; Cheeseman et al. 2001; Bouck and Bloom 2005; Thakur and Sanyal 2011). KT integrity can be affected by defect in individual KT ensemble. RNA-Seq data from the *nri1Δ/Δ nri2Δ/Δ* double mutant revealed a significant reduction in the levels of *NSL1* and *DAD4* transcripts which code proteins of central (Mtw1 sub-complex) and outer (Dam1 sub-complex) KT (Figure 2B), respectively (Cheeseman et al. 2002; Euskirchen 2002). As the alteration in the stoichiometry of the KT proteins within the sub-complexes is known to hamper KT integrity in *C. albicans* (Roy et al. 2011; Thakur and Sanyal 2012), we measured intensity of inner (Cse4-GFP), central (Mtw1-GFP) and outer (Dad2-GFP) proteins in the mutants to assess the KT integrity. We did not observe any defect in the integrity of the KT based on the analysis of the GFP intensity of these three fusion proteins (Figure S3A-F). Interestingly, at G2/M stage, unlike the wild type cells with characteristic bi-lobed GFP signal, a significantly higher population of cells exhibited mono-lobed GFP signal for all the three KT proteins in the *nri* mutants (Figure 5A-C). These results indicate that although the kinetochore integrity is not hampered in the *nri* mutants, the disjoining of sister KT clusters due to microtubule based pulling force, to take a bi-lobed organization is largely compromised. This might result from improper pulling force exerted by the microtubules that was found abnormal in the mutants and/or reduced stretchability of the centromere proximal chromatin in the mutants.

**Figure 5:**
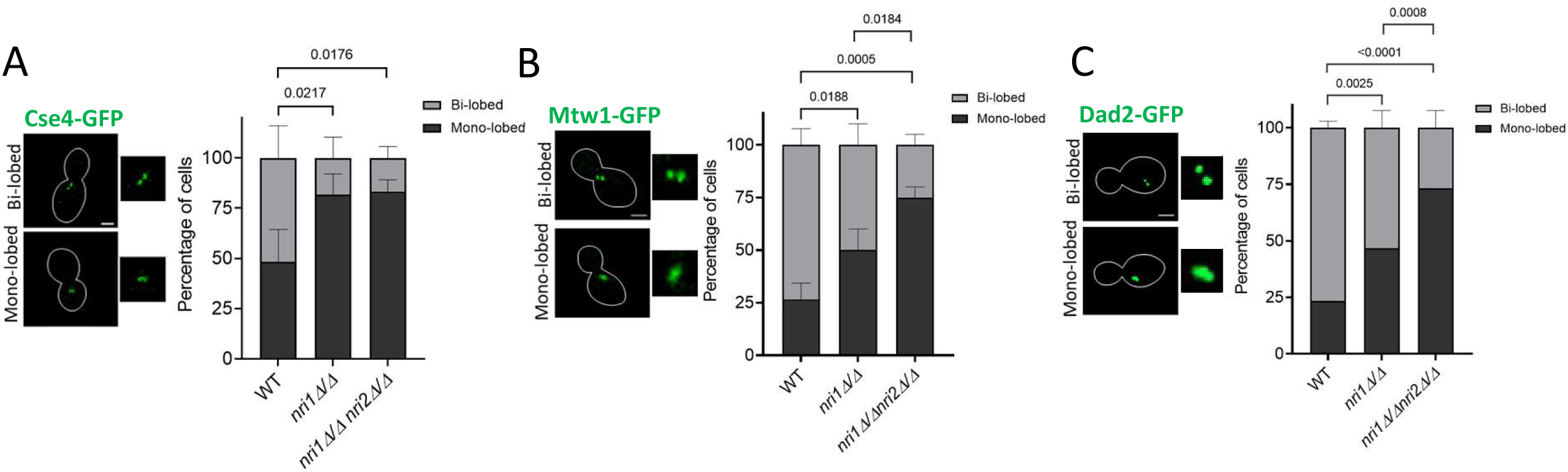
Bi-lobed organization of kinetochore clusters is affected in *nri1Δ/Δ nri2Δ/Δ* double mutant at G2/M stage. Left, representative images of the mono-lobed and bi-lobed signal types and right, their quantification in G2/M cell population harboring **A.** Cse4-GFP, **B.** Mtw1-GFP, and **C.** Dad2-GFP. Scale bar 2 µm. Statistical analysis was done by one-way ANOVA. Data from three biological replicates plotted with error bars indicating standard deviation. N = 30 for each cell cycle stage. Only significant p-value are mentioned in the graphs.

### Absence of *NRI* genes does not perturb centromeric cohesion but affects global chromatin

Various reports highlight the function of the RSC complex in establishment and maintenance of proper sister chromatid cohesion (Lopez-Serra et al. 2014; Muñoz et al. 2019; 2022). The correlation of proper cohesion and normal spindle structure is also deciphered in *S. pombe* and mice oocytes (Toyoda et al. 2002; Lu et al. 2018). Moreover, earlier we observed that cohesion and spindle mediated pulling forces determine the resolution of the *CEN7*-GFP signal (Sane et al. 2021). As we observed abnormal spindle morphology and significantly increased mono-lobed KT-GFP signal in the *nri* mutants, we evaluated the role of Nri proteins in the sister chromatid cohesion. For this, microtubules were depolymerized using 50 µg/ml nocodazole in *CEN7*::TetO tagged TetR-GFP expressing strains and nucleus was stained using live cell DAPI straining protocol (Sane et al. 2021). Depolymerization of the microtubules was confirmed by indirect immunofluorescence using anti-Tub1 antibody. No tubulin signal was observed in around 80% of the nocodazole treated cells for all the strains (Figure S4). GFP signal was observed in large budded cells with unsegregated DAPI to assess the centromeric cohesion in metaphase cells. In the cells with proper chromatid cohesion, two GFP signals from two sister chromatids coalesces into one signal (dot) due to the diffraction limit of the microscope (Figure 6A, type 1), whereas the signal is visible as two GFP dots (Figure 6A, type 2) in cells having defective cohesion. We found no significant difference in the type 1 and type 2 signal for the *nri* mutants as compared to the WT strain (Figure 6B, +NOC) indicating that sister chromatid cohesion is not perturbed in the mutants. In the cells with intact microtubules (-NOC), the sister chromatids experience outward pulling force by microtubules, and because of this, some population of cells harbor 2 GFP dots (Figure 6A, type 2) Notably, *nri* mutants displayed significantly less percentage of type 2 signal (Figure 6B, -NOC). To understand the reason behind reduction in the type 2 signal, we tagged Tub1-RFP for visualization of the spindles to assess the cell cycle stages, and quantified GFP signal only in the cells harboring metaphase spindle (0.5-2 µm length) (Figure 6C). At metaphase, 83% of the WT cells exhibited type 2 signal. However, 50% and 51% respectively of *nri1Δ/Δ* and *nri1Δ/Δ nri2Δ/Δ* mutant cells displayed significant reduction in type 2 signal (Figure 6D). This indicated that perhaps the force exerted by the spindles on the kinetochores in the *nri* mutants is not strong enough to resolve the GFP dots. The observed spindle morphology defects (Figure 4) in the mutants may account for this. Alternatively, but not mutually exclusive, the chromatin in the mutant may become too condensed to be separated by the spindle mediated opposite pulling forces. To test the condensation status of the chromatin, we examined its accessibility to micrococcal nuclease (MNase) in G2/M phase arrested WT and mutant cells and observed that in the mutant the chromatin is less accessible to MNase than in the wild type suggesting a more condensed nature of the chromatin in the mutant (Figure S5).

**Figure 6:**
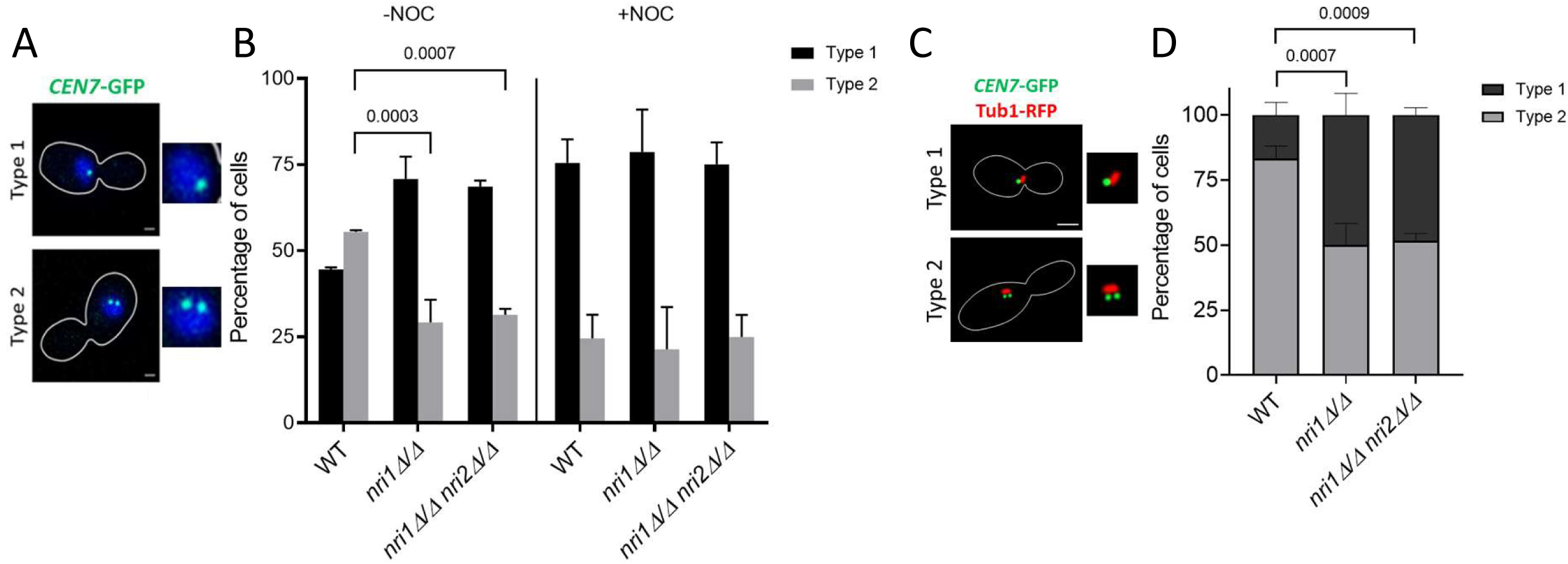
Absence of *nri* proteins does not affect sister chromatid cohesion. **A.** Representative images of two types of *CEN7*::TetO/TetR-GFP (*CEN7*-GFP) signals. Type 1, single GFP dot depicting cohesed sister chromatids and Type 2, two GFP dots depicting non-cohesed sister chromatids. Scale bar 2 µm. **B.** Bar graphs show percentages for the cells of the indicated strains treated with or without NOC. Error bar indicating standard deviation. Statistical analysis performed by ANOVA. Data from three biological replicates. N ≥ 99.**C.** Representative images of *CEN7*-GFP signal categories in metaphase cells showing 0.5-2 µm spindle (Tub1-RFP). Scale bar 2 µm. **D.** Quantification of the signal categories shown in C in the indicated cells. N ≥ 96. Data from three biological replicates. Error bars indicate standard deviation. Statistical analysis performed by ANOVA. Only significant p-value are mentioned in the graphs.

### Cells without the Nri proteins arrest at S phase

Although the above results indicate that the *nri* mutants harbor defects in kinetochore-microtubule related processes and thus arrest at G2/M in SAC dependent way, the possibility of an additional defect in S phase cannot be ruled out particularly when replication genes such as *ORC4* and *CDC45* were found as DEGs in the RNA-Seq data from the *nri* mutants (Figure 2B). Since the distinction between the S and G2/M phases using budding index analysis is not accurate as the bud size might increase in case of transient arrest, we used flow cytometry to judge the S phase progression proficiency of the mutant cells compared to the WT. We observed that a significant proportion of the *nri1Δ/Δ nri2Δ/Δ* cells showed S phase arrest (Figure 7A). After gating the population based on the WT peaks (Figure S6), we observed that 13% and 19.1% of the population was in the S phase for the WT and *nri1Δ/Δ* mutant, respectively. On the other hand, a much higher (40.8%) cell population was in the S phase for the *nri1Δ/Δ nri2Δ/Δ* double mutant. This resulted in concomitant reduction in the G1 and G2/M population in the double mutant. G1 population dropped to 17.8% as compared to 33.1% and 20.1% of the WT and *nri1Δ/Δ*, respectively. On the other hand, G2/M population decreased to 26.9% as compared to 40.3% and 47.6% in the WT and *nri1Δ/Δ*, respectively. Populations for all other cell cycle stages were comparable in all the strains (Figure 7B). FACS analysis suggests that at least a fraction of the large budded double mutant cells harboring unsegregated DAPI (from the budding index analysis) might still be in the S phase at arrested condition as they may harbor defects in DNA replication. Yeast cells with defective replication exhibit sensitivity to hydroxyurea (HU) and camptothecin (CPT), a topoisomerase I inhibitor (Nitiss and Wang 1988). We also observed that the *nri1Δ/Δ nri2Δ/Δ* mutant is sensitive to these drugs (Figure 7C-D) supporting their replication defects and resulting S phase arrest phenotype. We also tested the susceptibility of *nri* mutants to UV radiation. Both *nri1Δ/Δ* and *nri1Δ/Δ nri2Δ/Δ* mutants exhibited hypersensitivity to UV radiation (Figure 7E). However, we did not observe any sensitivity of *nri* mutants to another DNA damaging agent methyl methanesulfonate (MMS), indicating a possible involvement of Nri proteins in the regulation of only specific DNA damage repair pathways (Figure S7).

**Figure 7:**
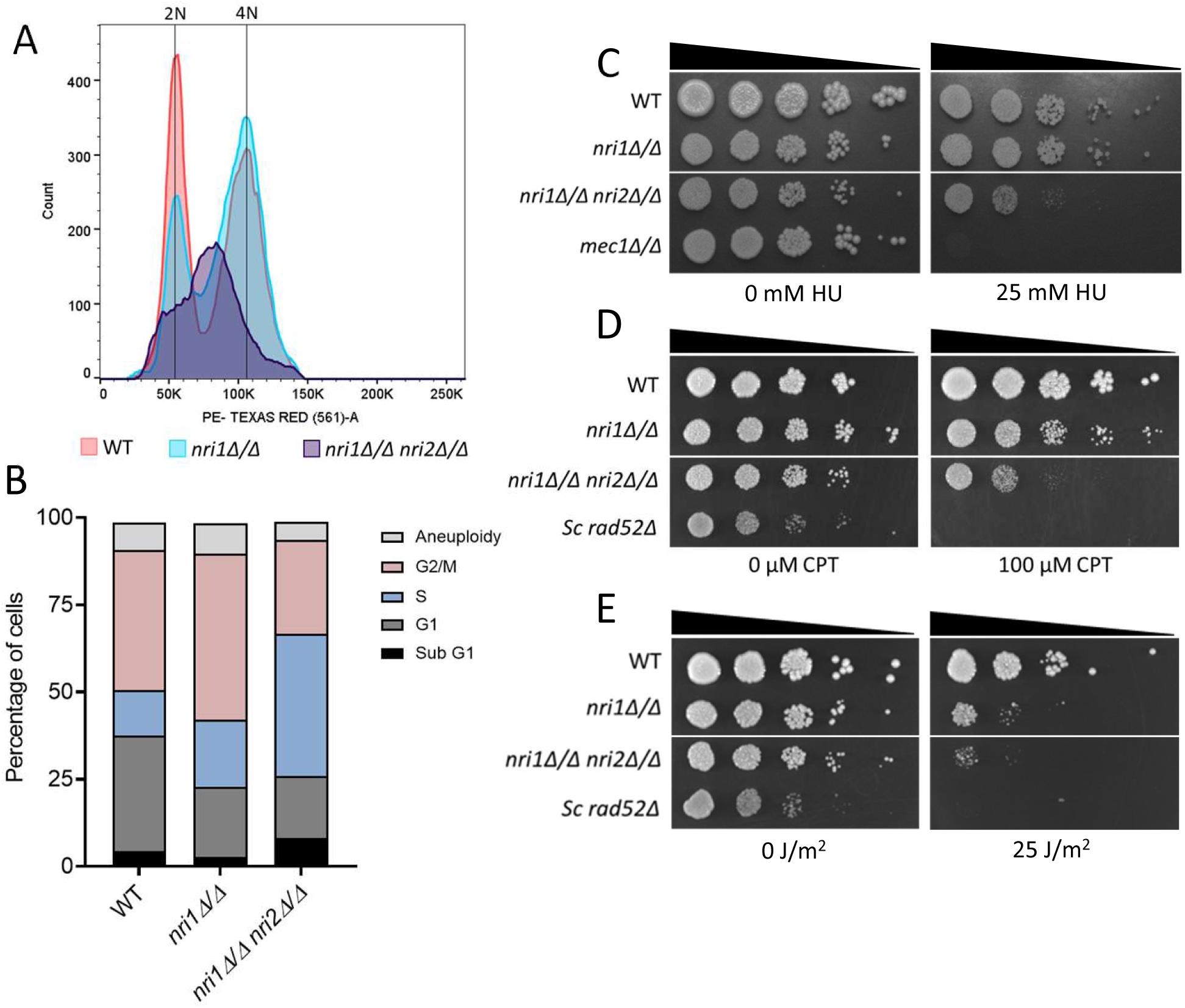
Cells lacking the Nri proteins show S phase arrest and drug sensitivity. **A.** Log phase cells were fixed with ethanol, treated with RNase, and isolated DNA was stained by propidium iodide. 20,000 events were recorded with BD FACS Aria flow cytometer with Texas Red filter. Gating was done using PI width (X-axis) vs PI area (Y-axis) to obtain single cell population. After gating, histograms of 18723, 18318, and 9899 events were plotted using PI area (X-axis) for WT, *nri1Δ/Δ*, and *nri1Δ/Δ nri2Δ/Δ* population, respectively. X-axis indicates PE Texas Red area (PI intensity) and Y-axis indicates cell count. WT cells showed two well separated peaks of 2N and 4N DNA content. While the *nri1* mutant showed similar two peaks like WT, the second peak in the double mutant largely remained in between 2N and 4N peaks. **B.** Quantification of the cells in different cell cycle stages based on the gating for PI peak intensity of the WT strain. Sensitivity was analyzed by spotting assay for **C.** HU, **D.** CPT, and **E.** UVC radiation. 10-fold serial dilutions of the cells of the indicated strains were spotted on YPDU plates containing indicated concentrations of the drugs or UVC radiation dose. Plates were incubated at 30°C for 48 hrs and imaged.

### Nri1 protein has the potential to activate transcription

The transcriptional activation exhibited by the chromatin remodeling complexes are often shown to be executed by one or more of their constituent proteins (Yoshinaga et al. 1992; Cosma et al. 1999; Yudkovsky et al. 1999; Johnson et al. 2008; Jian et al. 2021; Xue et al. 2021). As absence of Nri1 extensively affected *C. albicans* transcriptome, we wished to examine if it has transcription activation potential. To assess this, we evaluated sequence features of Nri1 using different bioinformatics tools (Figure 8A). Using PROSITE, we observed the presence of glutamine- and glutamic acid-rich stretches at the N- and C-terminal regions of Nri1 protein, respectively. Several reports suggest that glutamine-rich motifs play roles in transcription regulation (Ishii et al. 1997; Xiao and Jeang 1998; Sen et al. 2016). Additionally, the glutamic acid-rich regions within intrinsic disorder regions are shown to be involved in transcription activation (N 2013; Brodsky et al. 2020; Datta et al. 2025). Aligning with this, AIUPred (Erdős and Dosztányi 2024) also predicted that glutamic acid-rich region (residues 470-600) of Nri1 is disordered. Then, a motif search using Pfam database predicted transcription factor IIA, alpha/beta subunit motif between 109-219 amino acids of the Nri1 protein. To understand the evolutionary conservation of the amino acid residues, ConSurf analysis showed the N-terminal region (residues 1-92) is conserved across multiple proteins, including several predicted transcription factors from various *Candida* species (UniProt IDs: A0A8J5QLV9, H8WXN7, A0A8H7ZCI4, G8BF45, A0AAD5BGF6, A0A642UPY1). On the other hand, no domain/motif was predicted for Nri2 protein.

**Figure 8:**
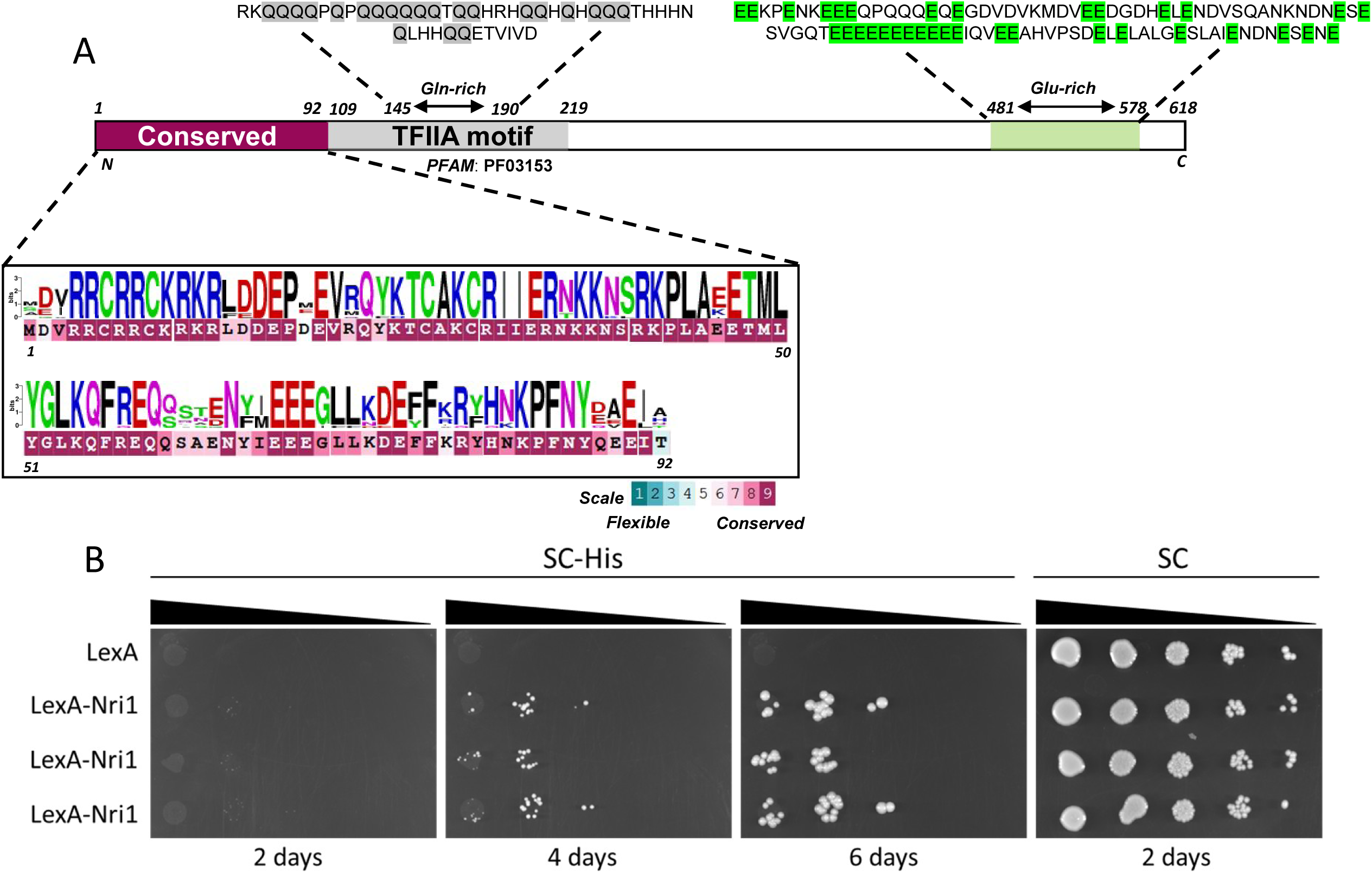
Nri1 protein exhibits a potential transcription activation ability. **A.** In-silico analysis revealed various features of the Nri1 protein. ConSurf analysis predicted 1-92 amino acids to be conserved, which displays high similarity with various transcription factors. ConSurf conservation scale ranges from 1 to 9, with highest conservation coloured in magenta. WebLogo was used to make the weighted sequence logo based on the ConSurf multiple sequence alignment. Motif search using Pfam database predicted 109-219 residues comprise transcription factor IIA (TFIIA) motif. PROSITE search detected 145-190 residues to be glutamine-rich (Gln-rich) and 481-578 residues as glutamic acid-rich (Glu-rich). **B.** *Candida* monohybrid assay detected the ability of the SC2H3 strain (5xLexO-*HIS1*) expressing LexA-Nri1, but not LexA alone, to grow on SC-His plate, indicating transcription activation potential of Nri1. Cells from three independent transformants were spotted on the SC-His and SC plates, incubated at 30 °C for the indicated time, and imaged.

We then adapted a CUG codon optimized mono-hybrid system to assess experimentally the transcription activation potential of the Nri1 protein (Stynen et al. 2010). If Nri1 has a transcription activation ability, a fusion protein of Nri1 and LexA DNA binding domain should be able to activate transcription of the reporter gene (*HIS1*) placed downstream of the LexA operator (5XLexO-*HIS1*), making the mono-hybrid strain (SC2H3) histidine prototroph. The LexA-Nri1 fusion protein was functional as it complemented the known stress sensitivity phenotype of the *nri1Δ/Δ* mutant (Figure S8). We observed that SC2H3 expressing LexA-Nri1, but not LexA alone, was able to grow on the histidine dropout plate, supporting the transcriptional activation potential of Nri1 protein (Figure 8B). However, further experiments are required to understand whether the observed transcription activation is achieved by Nri1 alone or in conjunction with other proteins of the RSC complex.

## Discussion

The RSC chromatin-remodeling complex is a multi-subunit complex that regulates myriad functions in eukaryotes (Clapier and Cairns 2009). Previous work by our group identified the composition of the *C. albicans* RSC complex and discovered that the complex harbors two novel CTG clade-specific proteins, namely Nri1 and Nri2 (Balachandra et al. 2020). As the species-specific subunits account for the observed functional variance of the complex across species and as there is a need to identify novel fungal-specific targets for the development of new anti-*Candida* drugs, in this study, we characterized the roles of the Nri proteins with respect to the cell cycle progression of the organism.

We observed that *NRI1* deletion or combined deletion of *NRI1* and *NRI2* resulted in a marked reduction in the growth rate of *C. albicans*, implying their significance in the cell proliferation (Figure 1A-C). On the other hand, *NRI2* deletion had no apparent growth defect (Figure 1A, S1A). Synthetic sick phenotype of the *nri1Δ/Δ nri2Δ/Δ* double mutant indicated that these genes might regulate cell proliferation via distinct pathways, with Nri1 playing a major role. This observed reduction in the growth rate occurred because of disruption in the cell cycle progression and not due to increased cell death (Figure S1C). Budding index analysis revealed that the cells lacking the Nri proteins tend to spend more time in G2/M and/or in S phase and additionally show cytokinesis defect as judged by increased percentage of multibudded cells (Figure 3A, S2). In *C. albicans*, impaired cell wall integrity is known to influence cytokinesis (Blankenship et al. 2010). Since the sensitivity of the *nri* mutants to cell wall-damaging agents indicates cell wall alteration (Figure S1B) (Dichtl et al. 2016), this can be a possible explanation for the observed defect in cytokinesis.

Since an accumulation of large budded cells with unsegregated DAPI in *nri1Δ/Δ nri2Δ/Δ* cells can account for defects both in G2/M and S phases, we investigated this further. We believe that there is a transient G2/M delay due to SAC activation as we observed genetic interaction between *NRI1* and *MAD2* genes (Figure 3B). Such delay was also reported for *rsc* mutants of ATPase subunit (Sth1) in *S. cerevisiae* (Tsuchiya et al. 1998; Hsu et al. 2003) and in *C. albicans* (Prasad et al. 2019). This argues that the Nri proteins are required for proper kinetochore-microtubule functions. But, interestingly, the effect of removing the Nri proteins is somewhat different from removing Sth1 of the RSC Complex. While both *nri* and *sth1* mutants showed microtubule depolymerizing drug sensitivity (Figure 4D, E; Hsu et al. 2003; Prasad et al. 2019), but unlike *sth1* mutant, *nri* mutants showed no sister chromatid cohesion (Figure 6A, B; Hsu et al. 2003; Prasad et al. 2019) but a massive anaphase spindle defect (Figure 4B; Hsu et al. 2003; Prasad et al. 2019). In budding yeast, RSC is shown to promote association of cohesin (Mcd1) with chromosomal arms but not with centromeres (Huang et al. 2004). Since with *CEN7*-GFP strain we could only detect centromeric cohesion, this can be one of the reasons why we did not observe a cohesion defect in the *nri* mutants.

In addition, surprisingly, we noticed non-disjunction of kinetochores or centromeric chromatin in a sizable fraction of the *nri* mutant cells (Figure 5, 6C, D) which was not reported for *sth1* mutant. The observed spindle defect (Figure 4) may account for this by not proving requisite pulling force, which needs to be measured. The non-disjunction may arise also due to higher compaction of the chromatin in the mutants (Figure S5) as the loss of RSC function is known to reduce DNA accessibility and hence to increase chromatin compaction by causing nucleosome crowding (Saha et al. 2005; Parnell et al. 2008; Chaban et al. 2008). As the increased chromatin compaction can resist splitting of the fluorescence signals, that might also explain the increased mono-lobed signal of the kinetochores or *CEN7*-GFP in the *nri* cells (Figure 5, 6D; Anbalagan et al. 2024). We observed variation in the mono- and bi-lobed percentages in *nri1Δ/Δ* and *nri1Δ/Δ nri2Δ/Δ* mutants for *CEN7*-GFP, Cse4-GFP, Mtw1-GFP, and Dad2-GFP signals. The difference in spatial distances of these signals from the centromere thus experiencing different microtubule pulling force may account for this variation. We also observed that the *nri1Δ/Δ nri2Δ/Δ* cells are impaired in timely completing the S phase using FACS analysis (Figure 7A, B) which is also reflected by their HU and CPT sensitivity (Figure 7C, D). Observed UV radiation sensitivity also indicated the possible roles of Nri proteins in the regulation of DNA damage repair mechanisms. This is not surprising given the fact that RSC, through nucleosome repositioning, facilitates replication by promoting repair of intrinsic DNA damage generated during replication (Shim et al. 2007; Niimi et al. 2012; Czaja et al. 2014).

Chromosome missegregation is a hallmark outcome of perturbation in chromatin, kinetochore or in microtubule spindle from yeast to humans (Skibbens and Hieter 1998; Malmanche et al. 2006). However, surprisingly, despite cell cycle progression defects and alterations in spindle morphology, kinetochore, and chromatin elasticity, we did not observe any gross chromosomal segregation defects in the *nri* mutants (Figure S1D). This is a remarkable divergence from frequently reported phenotypes for the mutants with defects in centromere, kinetochore or spindle (Pangilinan and Spencer 1996; He et al. 2001; Jones et al. 2001; Warren et al. 2002; Kiermaier et al. 2009; Parnell et al. 2024). The absence of such missegregation here might suggest that although cell cycle progression is perturbed in *nri* mutants, salvage mechanisms still sustain fundamental chromosome segregation fidelity and viability, at least under the examined conditions. Further study of cell cycle-dependent dynamics of chromosome segregation along with spindle behaviour in the *nri* mutants might provide mechanistic insights into how *C. albicans* can tolerate spindle and chromatin defects, potentially relevant for the adaptability and pathogenicity of the organism.

In summary, our results indicate that Nri proteins regulate *C. albicans* proliferation by controlling cell cycle progression at multiple stages. However, we cannot comment if Nri proteins perform any of their functions independent of the RSC complex. Additionally, the spectrum of DEGs indicates that the loss of Nri1 and Nri2 has widespread consequences, likely influencing various cellular pathways beyond the tested phenotypes in this study. The predicted TFIIA motif in Nri1 protein and evidence of LexA-Nri1 fusion protein activating transcription indicate potential direct and indirect roles of Nri1 in the regulation of gene expression. Notably, the presence of DEGs involved in the virulence-related biological processes also points towards the possible role of Nri proteins in *C. albicans* pathogenesis. As fungal fitness also affects its virulence, the potential role of Nri proteins, mainly Nri1, in the regulation of *C. albicans* virulence cannot be denied. Sensitivity of *nri* mutants to physiologically relevant stressors also supports this hypothesis (Figure S1B). Hence, a separate study is being carried out to delineate the pathogenic potential of the *nri* mutants. Furthermore, given the CTG clade-specific nature of the Nri proteins, a structural characterization of these proteins as physiological targets will be particularly beneficial to develop novel anti-*Candida* drugs.

## Acknowledgement

We acknowledge Prof. Judith Berman, Tel Aviv University, Israel and Prof. Kaustuv Sanyal, JNCASR, India for providing the Dad2-GFP strain. We acknowledge Prof. Patrick Van Dijck, Katholieke Universiteit Leuven, Belgium and Prof. Ambarish Kunwar, IIT Bombay, India for providing Candida optimized two-hybrid system and UV chamber, respectively. We thank Centre for Sophisticated Instruments and Facilities (CSIF) of IIT Bombay for NGS platform, Zeiss LSM780, Nikon confocal microscope and FIST grant (SR/FST/LSI-572/2013) from Govt of India for flow cytometer. SKG is supported by a grant (BT/PR45343/MED/29/1606/2022) from the Department of Biotechnology (DBT), Govt of India. AJ is supported by CSIR fellowship (09/087(1048)/2020-EMR-I), Govt of India. HK is supported by IIT Bombay IPDF fellowship. AS is supported by the New Faculty Seed Grant (NFSG/HYD/2023/H0866) from BITS-Pilani, Hyderabad Campus, and Core Research Grant (CRG/2023/006998) from Anusandhan National Research Foundation, Govt of India. GB is supported by Institute (Doctoral) fellowship from BITS-Pilani, Hyderabad Campus.

Author contributions: AJ and SKG conceptualized, designed experiments, analyzed and interpreted the data. AJ, SR, SS performed the experiments. HK, and AS performed in-silico sequence analysis of Nri proteins. GB and AS analyzed RNA-seq data. AJ and SKG wrote the manuscript. GB and AS edited and proofread the manuscript.

## Materials and Methods

### Strains, growth conditions and transformation

*C. albicans* strains and plasmids used in this study are mentioned in the supplementary tables S1 and S2. Primers used for strain construction and validation are mentioned in supplementary table S3.

All the *C. albicans* strains were grown in YPDU (1% yeast extract, 2% peptone, 2% dextrose, supplemented with 100 µg/ml uridine) medium at 30 °C, unless stated otherwise. Lithium acetate transformation protocol was used to construct *C. albicans* strains (Walther and Wendland 2003). For selection of transformants, YPDU+100 µg/ml nourseothricin or synthetic media without appropriate amino acid was used.

### Growth rate analysis

Overnight grown *C. albicans* culture was used to set OD_600_ of YPDU media to 0.15 and grown at 30 °C, 200 rpm. Thereafter, OD_600_ was measured every 1 hr till the culture reached saturation (15-16 hrs). Doubling time was calculated from the exponential growth phase.

### Spot dilution assay

10-fold serial dilutions of log phase WT and mutant strains were spot-inoculated in descending cell concentration based on the experiment: YPDU plate to evaluate growth defect at standard growth conditions; YPDU containing indicated concentrations of NOC and TBZ. Plates were incubated at 30 °C unless stated otherwise, and images were captured 1-2 days post-inoculation.

### Budding index analysis

Log phase cells were harvested and washed with 0.1 M phosphate buffer, pH 7.5 and permeabilized with 70% ethanol. The cells were again washed once with 0.1 M phosphate buffer, pH 7.5, and resuspended in 100 μl of 2 μg/ml DAPI solution. The tubes were incubated in the dark for 20 minutes, and the images were acquired using Zeiss Axio Observer Z1 microscope. Cells were classified based on the nuclear position and bud size.

### Zymolyase assay

Log phase cells were washed and resuspended in spheroplasting buffer. Cells were treated with 0.1 mg/ml zymolyase T20 (MP Biomedicals) for 15 mins, washed and observed under the Zeiss Axio Observer Z1. Percentage of multibudded cells was quantified for untreated and treated cells.

### DNA ploidy analysis

Flow cytometric analysis was performed according to Agarwal et al. 2015 with slight modifications. Briefly, 2 x 10^8^ log phase cells were harvested by centrifugation and washed once with 10 ml distilled water. Pellet was then resuspended in 100 µl D/W. Cells were fixed with 70% ethanol for 1 hr at RT in a tube rotator. Fixed cells were once washed with PBS and rehydrated by incubating in 1 ml PBS at 4 °C for 2 hrs. Rehydrated cells were then treated with 10 µg/ml RNase for 4 hrs at 37 °C. Followed by RNase treatment, cells were washed with PBS and overnight incubated in 4 °C. For propidium iodide staining, cells were incubated with 5 µg/ml PI solution at RT for 30 mins in dark. PI staining was assessed by Zeiss Axio Observer epifluorescence microscope. Cells were then diluted in a ratio of 1:4, vortexed, and sonicated with a probe sonicator for 10 seconds at 20% amplitude and immediately run in BD FACSAria™ Fusion flow cytometer with PE-Texas Red filter. Analysis was done using FlowJo software version 10.6.1 according to Todd et al., 2018. Parameters of detailed analysis are mentioned in the respective figure legends.

### Fluorescence imaging

Overnight grown cultures of fluorescently tagged WT and mutant strains were used to set the OD_600_ to 0.2 in fresh YPDU medium and cells were grown till OD_600_ 0.8-1. Cells were washed with 0.1 M phosphate buffer. Cse4-GFP and Mtw1-GFP samples were imaged with Zeiss L780 confocal microscope. Dad2-GFP samples were imaged with Nikon-Confocal AXR microscope.

### Sister chromatid cohesion (SCC) assay

SCC assay was performed according to Sane et al. 2021 with minor changes. Live cell DAPI staining protocol was followed to grow the cells till mid-log phase. At OD_600_ 0.4-0.5, culture was divided in two parts. Nocodazole to the final concentration of 50 µg/ml was added to one part and equal volume of DMSO (nocodazole solvent) was added to another part. Cells were grown at standard growth conditions for 2 hrs, washed twice with 0.1 M phosphate buffer and imaged. Indirect immunofluorescence for tubulin using anti-tubulin antibody (clone YOL1/34, Bio-Rad) was done to confirm microtubule depolymerization by nocodazole.

### RNA isolation

Total RNA was isolated from 4 x 10^7^ log phase cells. Cells were lysed by bead beating in 1 ml of RiboEx™ solution (GeneAll Biotechnology, South Korea). Lysate was centrifuged at 10,000 rpm for 1 min and supernatant was transferred in a fresh microcentrifuge tubes. 200 µl chloroform was added and mixed by inverting. Sample was incubated at RT for 2 mins and centrifuged at 10,000 rpm, 15 mins at 4 °C. Aqueous layer was transferred to a fresh tube and equal volume of isopropanol was added. Samples were incubated in -80 °C for 1 hr and centrifuged at 12,000 x g for 15 mins at 4 °C. Supernatant was discarded, pellet was washed with 75% ethanol, and air dried for 10 mins. Pellet was resuspended in 50 µl nuclease-free water. DNase treatment was done at 37 °C for 30 mins to remove any gDNA contamination. Followed by that, RNA was purified using RiboEx™ solution and chloroform, and precipitated using isopropanol. After washing with 75% ethanol and air drying, RNA pellet was finally resuspended in 30 µl NFW. RNA was quantified using NanoDrop microvolume spectrophotometer, and integrity was assessed by running 1 µg RNA on the agarose gel. PCR of 500 ng RNA template using *ACT1* qPCR primers was done to confirm the absence of gDNA in the samples. Isolated RNA was stored at -80 °C until further use.

### RNA-seq library preparation and sequencing

Before library preparation, isolated RNA was quantified using Qubit fluorimeter and RIN value was estimated by Agilent TapeStation. RNA-seq libraries were prepared in-house using the Illumina TruSeq Stranded Total RNA kit with 500 ng of total RNA (Illumina protocol 1000000040499 v00). Prepared libraries were quantified by Qubit fluorimeter and Agilent TapeStation. The library sizes ranged from 238-305 bp. 80-bp single-end sequencing of the libraries was done using Illumina NextSeq550 system generating approximately 36 million to 52 million reads per sample.

### RNA-seq data analysis

The fastq files containing the raw reads from RNA-seq were checked for quality using FastQC v0.12.1 (Andrews 2010; http://www.bioinformatics.babraham.ac.uk/projects/fastqc). Reads were trimmed using Trimmomatic v0.39 (Bolger et al. 2014) with the parameters ILLUMINACLIP:TruSeq3-SE.fa:2:30:10 LEADING:3 TRAILING:3 SLIDINGWINDOW:4:15 MINLEN:36. The trimmed reads were mapped to the *Candida albicans* SC5314, assembly 22 (A22) reference genome using HISAT2 v2.2.1 (Kim et al. 2019) using default parameters. For aligning RNA-seq reads, we used a modified FASTA file containing DNA sequences of one set (haplotype A) of homologous chromosomes (Ca22chr1A-7A, chrRA) and chrM from the phased, diploid A22 genome assembly of *C. albicans* SC5314 version_A22-s07-m01-r198_chromosomes.fasta [downloaded on 07-01-2024 from CGD]. The resultant BAM file was used along with the GFF file [C_albicans_SC5314_version_A22-s07-m01r198_features_with_chromosome_sequences.gff] downloaded on 07-01-2024 from CGD to obtain the count matrix containing the raw counts of reads mapped to haplotype A and mitochondrial transcripts for each sample using the Subread package (featureCounts v2.0.6) (Liao et al. 2014). The ‘-g Parent’ parameter was specifically set to count the reads mapped to the transcripts in the GFF file. The count matrix file was used to identify the differentially expressed genes (DEGs) using DESeq2 v1.42.1 package (Love et al. 2014) of R v4.3 with cutoff criteria of log2FoldChange of >1 or <-1, and FDR (adjusted p-value) of <0.05. The associated data for *nri1Δ/Δ* single mutant and *nri1Δ/Δ nri2Δ/Δ* double mutant w.r.t. WT are provided in the Supplementary file. CGD GO slim mapper tool was used for Gene Ontology analysis.

### In-silico analysis of Nri proteins

The protein sequences of Nri1/2 were obtained from the CGD. Further, these protein sequences were used for the detection of protein domain/motif/family using Motif Search (https://www.genome.jp/tools/motif/) that scans them against a collection of motifs in the Pfam database (Mistry et al. 2021), as well as using PROSITE Scanning (Sigrist et al. 2013). Further, the ConSurf web server (Yariv et al. 2023) was used to identify functionally important and conserved regions in the proteins. The proteins obtained from the ConSurf server were further aligned using the CLUSTALW (Thompson et al. 1994), and the weighted sequence logo was obtained using the WebLogo server (Crooks et al. 2004).

## Data availability

Output files for RNA-seq analysis are uploaded as supplementary information. RNA-seq raw data will be deposited to the Gene Expression Omnibus (GEO) repository before the final publication. However, data can be made available to the reviewers on demand.

